# A Meta-model for ADMET Property Prediction Analysis

**DOI:** 10.1101/2023.12.05.570279

**Authors:** Sarala Padi, Antonio Cardone, Ram D. Sriram

## Abstract

In drug discovery analysis chemical absorption, distribution, metabolism, excretion, and toxicity (ADMET) properties play a critical role. These properties allow the quantitative evaluation of a designed drug’s efficacy. Several machine learning models have been designed for the prediction of ADMET properties. However, no single method seems to enable the accurate prediction of these properties. In this paper, we build a meta-model that learns the best possible way to combine the scores from multiple heterogeneous machine learning models to effectively predict the ADMET properties. We evaluate the performance of our proposed model against the Therapeutics Data Commons (TDC) ADMET benchmark dataset. The proposed meta-model outperforms state-of-the-art methods such as XGBoost in the TDC leaderboard, and it ranks first in five and in the top three positions for fifteen out of twenty-two prediction tasks.

## 1 Introduction

In the context of drug discovery, the in-silico characterization of quantitative structureactivity relationships (QSAR) is of key importance for building useful models. Also, the pharmacological activity of a chemical compound is described based on ADMET properties. Over the past decade, pharmaceutical and biotech companies have invested heavily in high accuracy testing and measurement capabilities, which have led the experimental characterization of millions of compounds and their associated ADMET properties. In order to take advantage of the widespread availability of QSAR and ADMET data and tackle its inherent complexity, scientists have resorted to machine learning for in-silico modeling of optimal drug leads. Successful modeling efforts have reduced the need for in-vitro or in-vivo experiments and have led to faster identification of promising drug compounds [1–3].

There are various molecular representations that can be used for cheminformatics, but they often involve some level of abstraction and loss of information. Designing machine learning approaches for ADMET prediction is also difficult because of the presence of complex non-linearity in the training data and of the difficulty in selecting optimal parameter sets. However, the random forest approach has shown promising performance in drug design when combined with specialized molecular fingerprints, and it is robust with respect to the associated hyper-parameters [4–8].

Deep learning models are becoming increasingly popular in predicting ADMET properties. However, the process of building these models can be challenging due to the limited amount of training data available, especially when human subjects are involved. Additionally, it is difficult to obtain balanced data, and there are often issues with variability, imbalance, quality, and missing values in the published data [9]. Another major challenge is the lack of explainability or interpretability in deep learning models used in biomedical science. It is important to understand the inner workings of the employed Artificial Intelligence (AI) model to make accurate interpretations of AI-driven predictions. There are ongoing efforts to address this challenge and gain a better understanding of how AI models work [10]. Unfortunately, practitioners are sometimes forced to rely on drug property predictions without fully understanding the criteria behind such predictions [11–13].

As we delve into machine learning applied to drug property prediction, we discover that several models have been developed to predict the ADMET and QSAR properties [14]. The commonly used algorithms include K-nearest neighbors and Support Vector Machines. Although these traditional models are user-friendly, they are sensitive to outliers and datasets that are not balanced. To address this challenge, tree-based algorithms have been used. However, noise in the data can lead to overfitting, making it challenging to fine-tune the hyper parameters of a given model [9]. In ADMET property prediction, usually a large number of different properties need to be predicted. Building and fine-tuning different models for each property prediction task can be difficult. It is not possible to build one model that can predict all different properties of ADMET or QSAR. This is where ensemble learning models come into play.

Ensemble learning is a popular machine learning approach, which combines prediction scores from multiple weak learners to enhance the final prediction performance of the model [15]. The ensemble learning methods include bagging, stacking, and boosting. Bagging involves training base models by taking into account a subset of data samples from the given set. For instance, random forests combines predictions from decision trees to generate the final predictions [16]. On the other hand, boosting learns from its previous weak learners’ errors to create a better prediction model. Two popular boosting models are XGBoost and AdaBoost [17]. The primary difference between boosting and bagging is that the former considers the entire dataset to build weak learners, while the latter uses only a subset of data to construct its base learners. On the other hand, the stacked model, which differs from bagging and boosting methods in that it utilizes multiple heterogeneous models to gain insight [18, 19]. Bagging and boosting methods rely on weak, homogeneous learners, whereas a stacked model combines the predictions of heterogeneous weak learners using a meta-learner to make better predictions [20].

In this paper, we explore a meta-model that combines the scores from multiple models to improve the model’s generalization. We show that our meta-model performs better than bagging and boosting-based models for ADMET prediction tasks. Our method also outperforms state-of-the-art (SOTA) in the Therapeutics Data Commons (TDC) ADMET prediction leaderboard, where meta-model ranks first in five and is in the top three positions for 15 out of 22 prediction tasks. Additionally, we show that combining heterogeneous models produces better prediction results than the state-of-the-art XGBoost model [21], leading to improved ADMET property prediction performance.

## 2 Related Work

Machine Learning is important for solving problems in areas like drug design, medical imaging, and drug discovery[22–24]. There are, however, specific challenges when applying these models to drug discovery applications. As mentioned above, one such challenge is the representation of data. The Simplified Molecular Input Line Entry Specifications (SMILES) string is a widely used molecular representation in molecular design. It is a linear form in which strings represent the molecular structure as a sequence of characters [25]. Each string contains the whole molecular structure, including identifiers for atoms and identifiers denoting topological features, such as bonds, rings, and branches. SMILES strings are either directly fed to the ML models or converted into feature-based representations, such as molecular fingerprints, one hot encoding, word embeddings, or even graph representations[26–29].

A variety of approaches for drug property prediction can be found in the literature, with specific focus on ADMET properties. DeepPurpose is a deep network that takes SMILES as input for ADMET property prediction [30]. It comprises an encoder that generates embeddings for the input representations, and a subsequent decoder that produces the property prediction output. In the DeepPurpose framework, over 50 state-of-the-art deep learning models are used to predict the drug properties. The two prominent issues with deep learning-based models are the need for a large amount of training data and the lack of interpretability, which may cause distrust in the biodemdical field. The AttentiveFP employs a graph-based attention mechanism, which enhances its predictive performance due to its ability to learn intermolecular and intramolecular interactions [31]. This method is more interpretable than deep learning models. In particular, the insight into molecular interactions leads to better drug property prediction accuracy. However, graph neural networks have not yet been fully assessed in the biomedical field.

Another class of models employing gradient-boosting techniques, such as XGBoost, relies on the sequential training of a series of decision trees. It uses a regularization term to reduce the overfitting of specialized fingerprints and descriptors representing molecular features of interest [17]. The XGBoost model achieved state-of-the-art accuracy for AMDET prediction analysis and other machine learning-based applications [21].

## 3 Meta-model

Ensemble learning is a powerful technique that combines the predictions of multiple models [32, 33]. However, a common drawback of this approach is that each model contributes equally to the ensemble predictions regardless of their performance. To address this issue, we can use a weighted average ensemble approach, which gives more weight to the high-performing models leading to significant improvements in model performance. Stacked generalization, known as meta-learning, takes this approach to the next level by replacing the linear weighted sum used to combine the submodels’ predictions with a learning algorithm [20]. The meta-learner can learn to effectively combine the scores from heterogeneous models, leading to even better predictions. Through this two-level learning process (Figure 1), models at the first level learn to make predictions from the given data, while at the second level the meta-learner learns to combine these scores and make predictions accordingly. This cutting-edge technology allows us to attain even greater accuracy and improves the model generalization for multiple tasks.

**Fig. 1.**
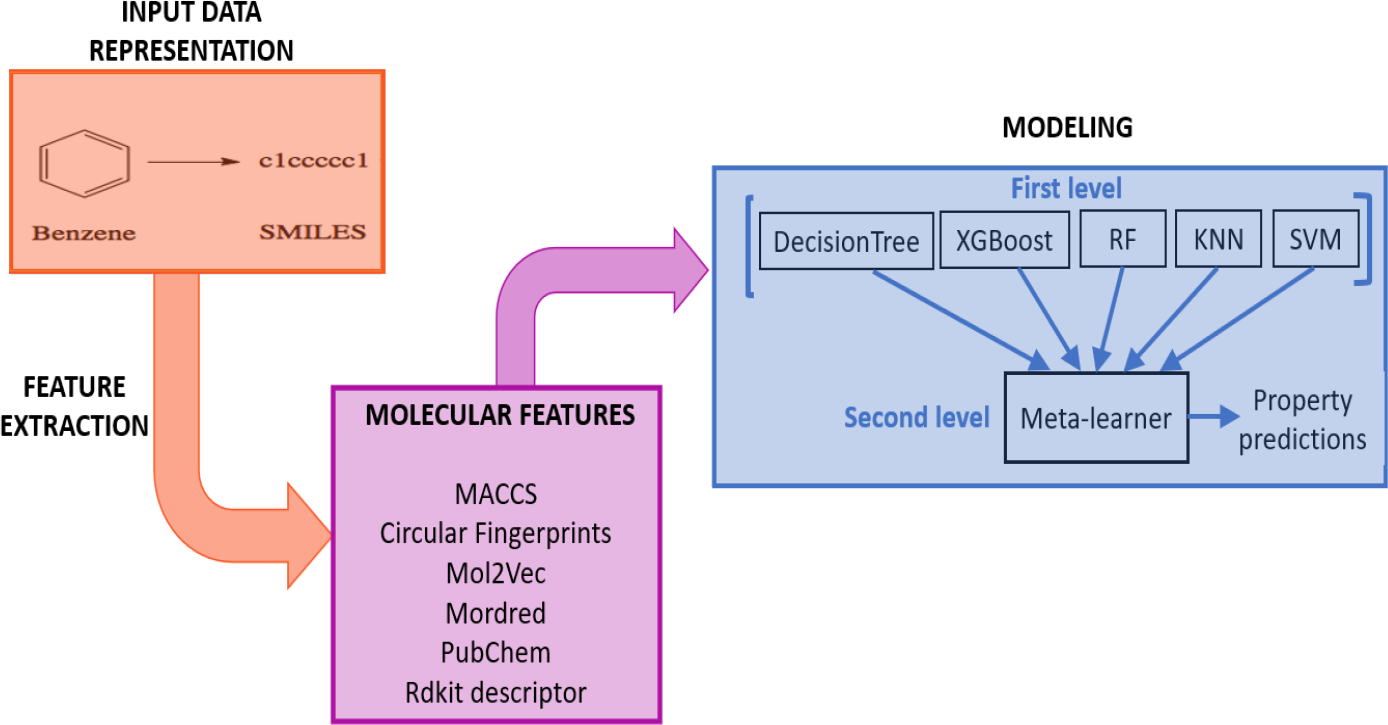
A meta-model used for ADMET property prediction analysis. Abbrev: SVM:Support vector machine, RF: Random forest,KNN:K-Nearest neighbour

The methodology used for ADMET property prediction analysis is illustrated in Figure 1. The machine learning model in the methodology is based on extracted feature descriptors for SMILE sequences using RDkit. At the first level, the methodology comprises of five machine learning models, including Support Vector Machines, Random Forest, XGBoost, K-nearest neighbors, and Decision Trees, which are heterogeneous models, along with a linear prediction model (second level) as the meta-learner. The meta-learner is responsible for learning how to combine the scores from the different models to improve prediction accuracy and gain a better understanding of the underlying data.

## 3.1 Fingerprints and descriptors

The molecular descriptors are a means of expressing the chemical properties of a given molecule. Molecular fingerprints are commonly used to generate molecular representations, as they encode the representation of structural or functional descriptors of molecules in a string format. For example, circular fingerprints, especially extended-connectivity fingerprints, are used to build machine-learning methods for modeling QSAR for biological analysis[34, 35]. To compute fingerprints and descriptors, we utilize the following six features from DeepChem [36]^1^:

- The MACCS fingerprints are commonly used structural keys that compute binary strings from the molecular structure.
- Circular fingerprints with extended connectivity are used to model structural activity by breaking up a molecule into circular neighborhoods.
- Mol2Vec[37] fingerprint is a vector representation of a molecules generated by an unsupervised machine learning approach.
- PubChem fingerprint contains 881 structural keys covering a wide range of substructures and features, and they are used in PubChem for similarity searching.
- Mordred descriptors [38] are used to calculate a set of chemical descriptors, such as the count of aromatic atoms or the count of halogen atoms.
- RDKit descriptors calculate a set of chemical descriptors, like the molecular weight and the number of radical electrons.

## 4 Dataset

### 4.2 Therapeutics Data Commons (TDC)

The Therapeutics Data Commons (TDC) [39] is the first unifying platform for systematically assessing and evaluating machine learning models across the entire range of therapeutics. It consists of 66 datasets and 22 learning tasks aimed at finding safe and effective medicines. TDC provides tools and resources, such as data functions, data splits, measures for model evaluation, and molecule generation tools. For each ADMET prediction task, TDC divides the dataset into predefined sets, with 80% of data for training and 20% of data for testing using a scaffold split mechanism. The scaffold split mechanism allows a machine learning model to predict the ADMET properties of drugs that are structurally different. The ADMET benchmark dataset is freely available for research^2^. Table 1 shows the ADMET dataset, the type of ML models built for each task, the metrics used to evaluate the models, and the number of data samples used to train and evaluate ML models.

**Table 1.**
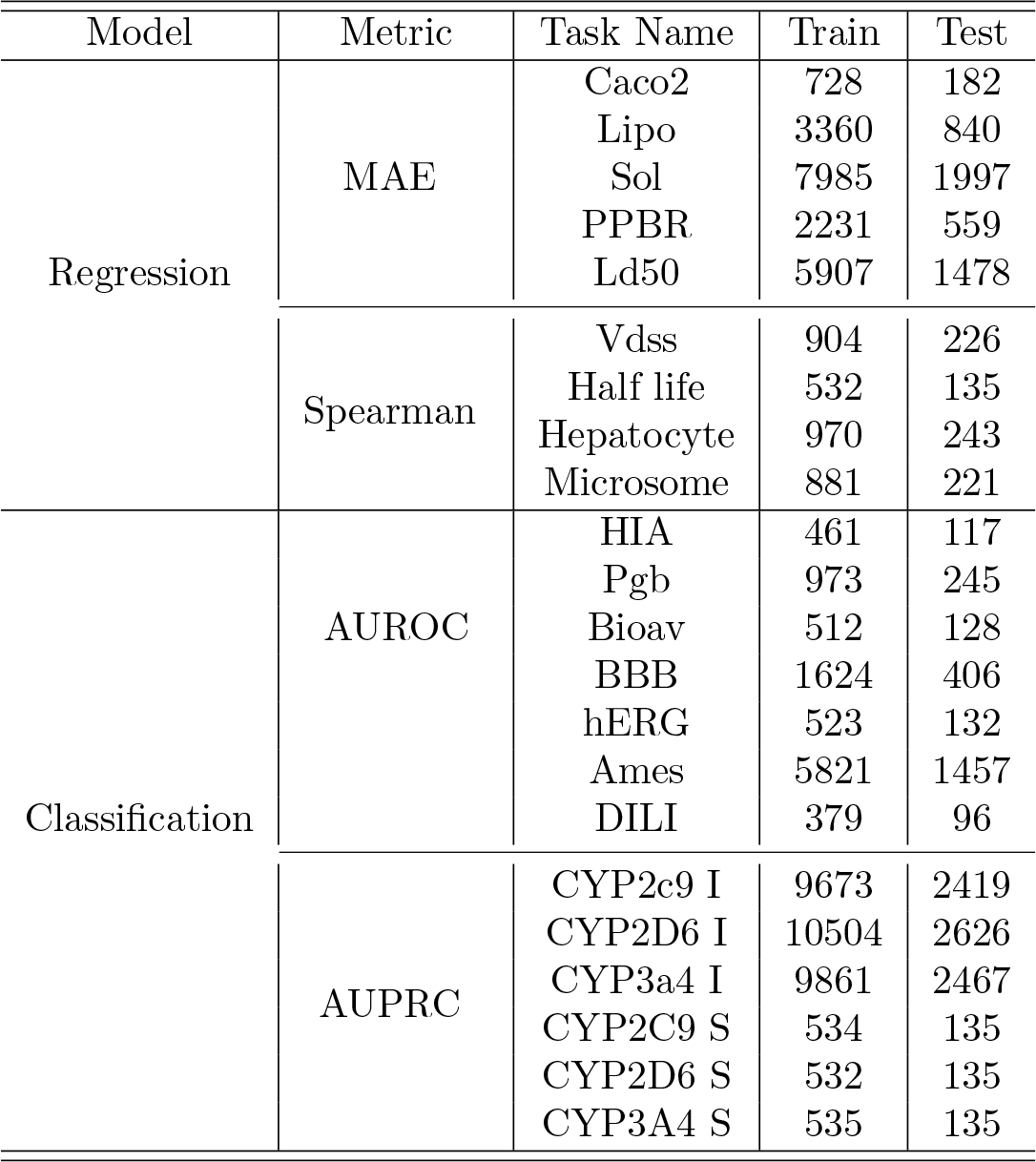
The table provides an overview of experimental settings used for ADMET property prediction analysis which includes the type of machine learning model used, associated metrics for evaluating model performance, and the number of examples used to build and test the models.

## 5 Experimental Setup

To implement the meta model, we use the Scikit-learn package which is available at^3^. The meta model combines estimators to reduce their biases[20]. The model predictions from the estimator are stacked together and used as input to a final estimator for the purpose of calculating prediction scores using a cross-validation strategy.

Table 1 shows that there are 22 tasks in ADMET prediction, of which nine are regression tasks, and 16 are binary classification tasks. The metrics for regression tasks are the mean absolute error (MAE) and Spearman correlation coefficient. We report the average receiver operating characteristics (AUROC) and the average precision and recall (AUPRC) metrics for the classification task.

A five-fold cross-validation method is used to train our meta-model. Hao et.al. [21] performed a grid search to fine-tune the parameters of the XGBoost model. But for our meta-model, the search space is big, and it is expensive to fine-tune all the parameters using a random grid search. Furthermore, it may not always result in the best global parameters. Thus, we perform our experiments using the default parameter settings (Table 2) except for XGBoost model^4^. The source code for our meta-model ia available at gitlab repository^5^.

**Table 2.**
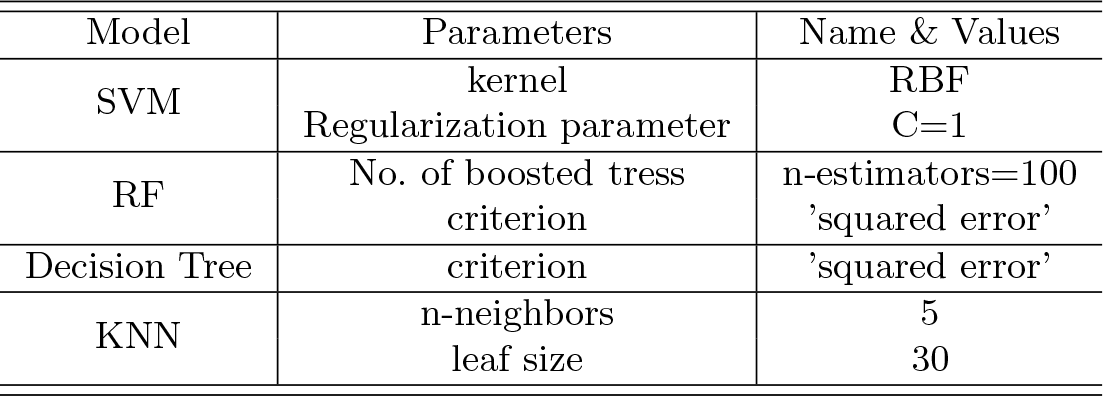
The parameter settings used for each model in the meta-model. Abbrev: SVM: Support vector machines; RF: Random forest classifier; KNN: K-nearest neighbours; RBF: Radial basis function kernel

| Model | Parameters | Name & Values |
| --- | --- | --- |
| SVM | kernel | RBF |
|  | Regularization parameter | C=1 |
| RF | No. of boosted trees | n-estimators=100 |
|  | criterion | 'squared error' |
| Decision Tree | criterion | 'squared error' |
| KNN | n-neighbors | 5 |
|  | leaf size | 30 |

## 6 Results

As part of the first step, we trained five models, including SVM, RF, XGBoost, KNN, and Decision Tree, and evaluated these models for ADMET prediction analysis. Later, we combined these model scores along with the meta-learner, linear regression, to further improve the generalization performance across ADMET tasks. The Figure 2 shows the comparison of five model performance’s with the meta-model. From the figure, we can see that the meta-model outperforms the other models in all the tasks. The meta-model achieves superior performance relative to individual classifiers due to a strong correlation between the various classifiers. This dramatically increases the meta-modal performance. On the other hand, the meta-model performs poorly for the “Half-Life” task because of a disagreement between the models. Note that the models are evaluated by considering random parameters. However, it would be beneficial to explore the parameter space of individual models in order to further enhance meta-model performance.

**Fig. 2.**
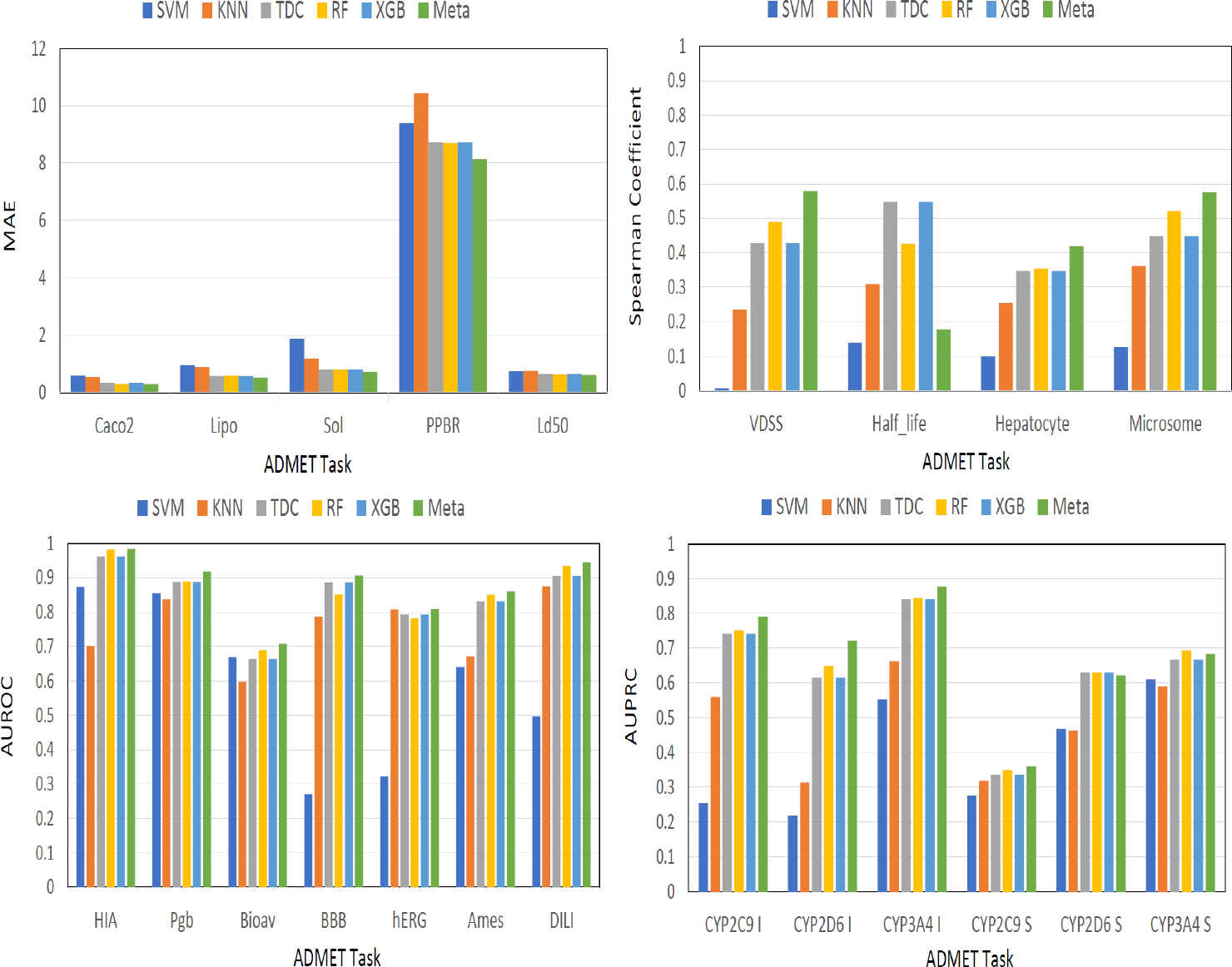
Performance comparison of the proposed meta-model with state-of-the-art models for the ADMET property prediction tasks.

In our study, we also conducted an experimental evaluation to compare the performance of the meta-model against the SOTA (Table 3) method in the TDC leaderboard for the ADMET prediction task. After analyzing the best-performing model for each task from the TDC leaderboard, we compared it to our meta-model performance. The results, as shown in Table 3, indicate that our model ranked first for five tasks and in the top three positions for 15 out of 22 tasks. These findings suggest that incorporating heterogeneous models can result in improved performance, and our meta-model effectively addresses the complexity involved in selecting a model and its parameters for each prediction.

**Table 3.** Evaluation of the meta-model for the ADMET property prediction and comparison with the state-of-the-art methods (SOTA) in the TDC leaderboard. Note: In the leaderboard, the top 3 positions of the meta-model are highlighted in bold.Abbrev: BB: BaseBoosting, RF: Random Forest

| Property | TDC |  | SOTA |  | Ours |  |
| --- | --- | --- | --- | --- | --- | --- |
|  | Task Name | Metric | Method | Score | Score | Rank |
| Absorption | Caco2 | MAE | BB | 0.285 ± 0.005 | 0.296 ± 0.011 | **3** |
| | HIA | AUROC | RFStacker | 0.988 ± 0.002 | 0.985 ± 0.002 | **3** |
| | Pgp | AUROC | BB | 0.946 ± 0.001 | 0.918 ± 0.001 | 6 |
| | Bioav | AUROC | SimGCN | 0.748 ± 0.033 | 0.709 ± 0.005 | **2** |
| | Lipo | MAE | XGBoost | 0.533 ± 0.005 | 0.521 ± 0.004 | **1** |
| | AqSol | MAE | XGBoost | 0.727 ± 0.004 | 0.731 ± 0.003 | **2** |
| Distribution | BBB | AUROC | BB | 0.923 ± 0.002 | 0.906 ± 0.004 | **2** |
| | PPBR | MAE | XGBoost | 8.251 ± 0.115 | 8.148 ± 0.172 | **1** |
| | VDss | Spearman | XGBoost | 0.612 ± 0.018 | 0.579 ± 0.023 | **3** |
| Metabolism | CYP2C9 Inhibition | AUPRC | XGBoost | 0.794 ± 0.004 | 0.791 ± 0.003 | **3** |
| | CYP2D6 Inhibition | AUPRC | BB | 0.721 ± 0.001 | 0.721 ± 0.003 | **1** |
| | CYP3A4 Inhibition | AUPRC | BB | 0.882 ± 0.001 | 0.877 ± 0.002 | **2** |
| | CYP2C9 Substrate | AUPRC | RF | 0.437 ± 0.022 | 0.36 ± 0.022 | 8 |
| | CYP2D6 Substrate | AUPRC | BB | 0.711 ± 0.006 | 0.622 ± 0.022 | 5 |
| | CYP3A4 Substrate | AUPRC | XGBoost | 0.680 ± 0.005 | 0.683 ± 0.004 | **1** |
| Excretion | Half Life | Spearman | BB | 0.416 ± 0.009 | 0.177 ± 0.08 | 8 |
| | CL-Hepa | Spearman | BB | 0.491 ± 0.006 | 0.419 ± 0.013 | 4 |
| | CL-Micro | Spearman | RFStacker | 0.625 ± 0.002 | 0.576 ± 0.007 | 7 |
| Toxicity | LD50 | MAE | autoML | 0.588 ± 0.005 | 0.605 ± 0.003 | **3** |
| | hERG | AUROC | RF | 0.875 ± 0.003 | 0.809 ± 0.008 | 4 |
| | Ames | AUROC | BB | 0.865 ± 0.002 | 0.861 ± 0.002 | **2** |
| | DILI | AUROC | BB | 0.937 ± 0.004 | 0.945 ± 0.006 | **1** |

A limitation of the meta-model is the associated inability to interpret the model or determine which features contributed the most to the prediction outcome. In order to address this issue, we decided to examine the feature importance scores of the XGBoost and Random Forest models to see how they compared. We consider XGBoost and Random Forest models because these two models are part of our meta-model and are proven to be good perfroaming models for ADMET prediction tasks. Figure 3 showcases the box plots of the feature importance scores for both models. Upon analyzing the XGBoost model, we found that the “Mordred” feature played a crucial role in predicting ADMET property, accounting for 60% of the contribution. On the other hand, the same feature contributed 70% in the Random Forest model. It is important to note that the significance and impact of a particular feature on prediction analysis vary depending on the prediction model employed.

**Fig. 3.**
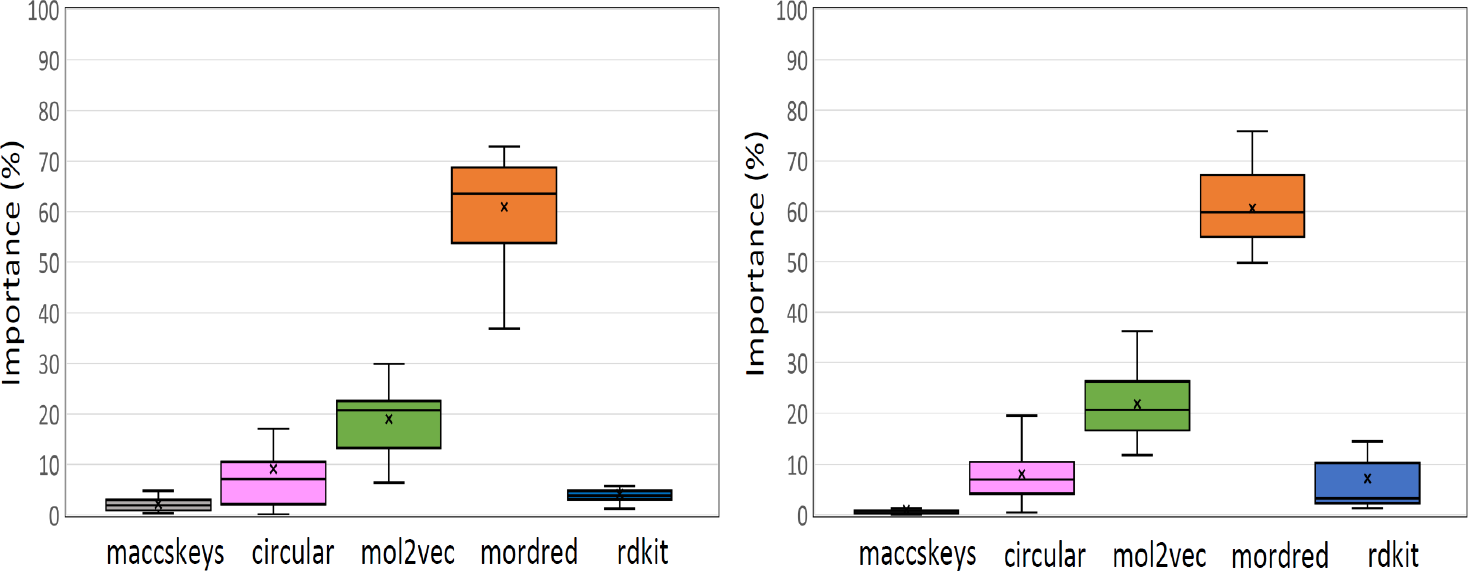
The feature importance of XGBoost (Left) and Random Forest (Right) models for the ADMET property prediction tasks.

## 7 Conclusion

This paper study the performance of a meta-model that combines multiple heterogeneous machine learning models for the ADMET property prediction analysis. We evaluated the meta-model performance on the TDC ADMET benchmark dataset for 22 different tasks. Our model ranked first for 5 tasks and in the top 3 positions for 15 out of 22 tasks in the TDC leaderboard. Our results demonstrate that the proposed meta-model outperforms five machine learning models, including state-of-the-art XGBoost model. This shows that the combination of bagging and boosting-based models provides additional and complementary insight, thereby increasing the prediction performance.

## Acknowledgements

The authors would like to thank Marcin Kociolek and Michael Majurski for their helpful comments and suggestions for the paper.

## Disclaimer

Certain equipment, instruments, software, or materials are identified in this paper in order to specify the experimental procedure adequately. Such identification is not intended to imply recommendation or endorsement of any product or service by NIST, nor is it intended to imply that the materials or equipment identified are necessarily the best available for the purpose.

## Footnotes

1 We utilize the features provided by [21]

2 https://tdcommons.ai/benchmark/admetgroup/overview/

3 https://scikit-learn.org/stable/

4 We set the XGBoost model parameters to be similar to[21]

5 https://gitlab.com/pscolab4/ADMET.git

